# Bioactive 3D-bioprinted scaffolds with endothelial progenitor cells and Zn^2+^-enhanced SFMA/HisMA/nHAP for healing of infected bone defect

**DOI:** 10.1101/2025.09.15.676384

**Authors:** Yunliang Zhu, Jiazhao Yang, Kai Xie, Jinke Chang, Shiyuan Fang, Jinsen Lu

## Abstract

Infected bone defects remain one of the greatest challenges in orthopedics, as bacterial contamination and insufficient vascularization severely compromise regeneration. Here, we report the development of a multifunctional 3D-bioprinted scaffold composed of zinc ion (Zn²□) functionalized silk fibroin methacryloyl (SFMA), histidine methacryloyl (HisMA), and nano-hydroxyapatite (nHAP), further loaded with endothelial progenitor cells (EPCs) to promote simultaneous antibacterial, angiogenic, and osteogenic responses. The photocrosslinkable SFMA/HisMA bio-ink reinforced with nHAP provided mechanical stability, controlled swelling and degradation, while Zn²□ coordination endowed strong antibacterial activity against E. coli and S. aureus. EPCs adhered and proliferated on the scaffolds, and in co-culture with bone marrow stromal cells (BMSCs) markedly enhanced osteogenic differentiation, as evidenced by increased expression of OCN, RUNX2, COL1, and OPN. Proteomic profiling further revealed that EPC-loaded scaffolds modulated osteoblast protein expression patterns consistent with both immunoregulation and active extracellular matrix reconstruction to promote tissue regeneration. In a murine femoral infected defect model, EPC-loaded scaffolds suppressed pro-inflammatory cytokines (IL-6, TNF-α), stimulated angiogenesis, and supported robust new bone formation, leading to accelerated defect repair confirmed by µCT and histological analyses. Together, these findings demonstrate that the EPC-loaded Zn²□ functionalized SFMA/HisMA/nHAP scaffold integrates antibacterial defence with vascular and osteogenic stimulation, offering a promising translational strategy for the treatment of complex infected bone defects.

**Graphic Abstract:** 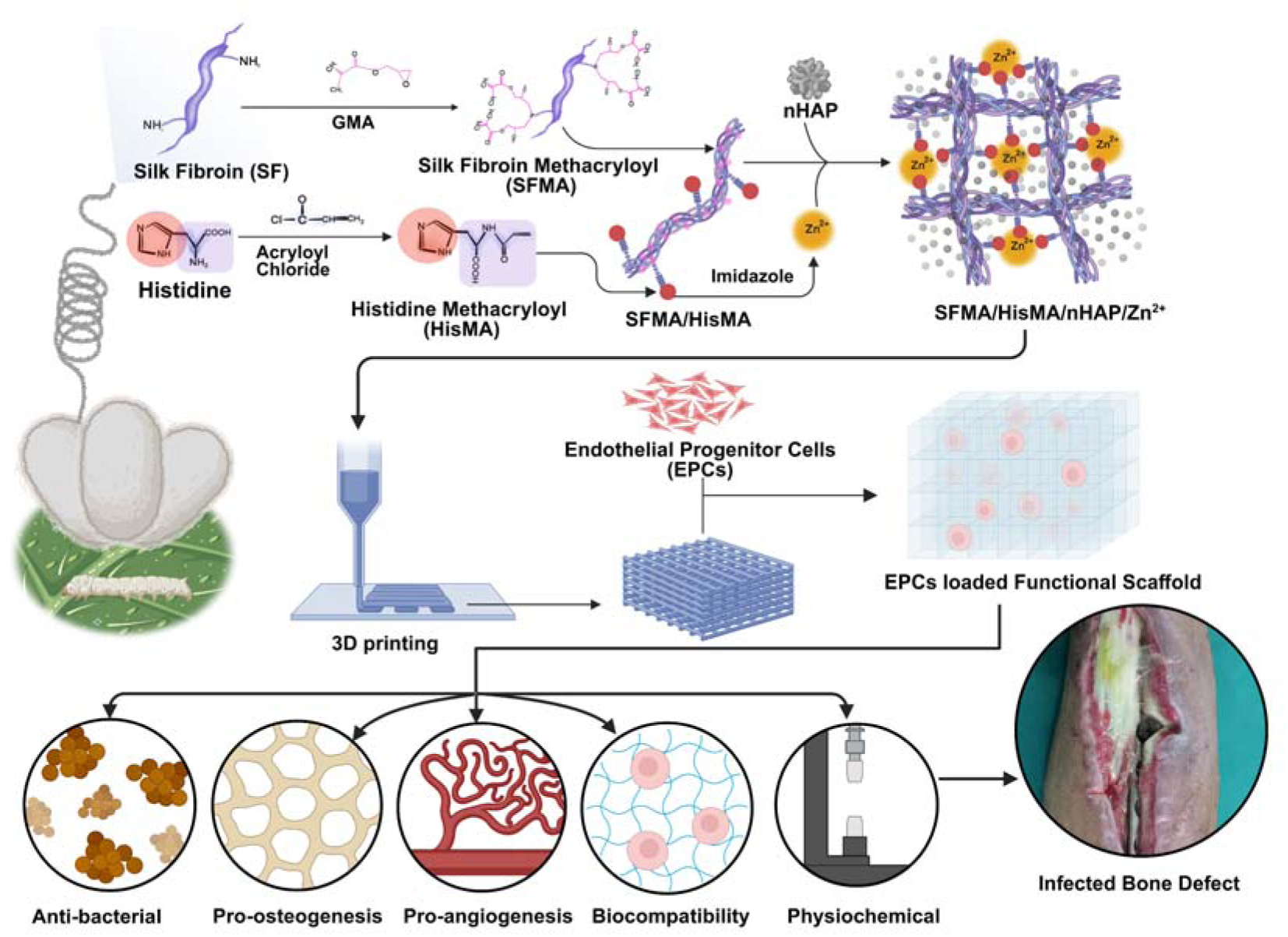

Schematic illustration of materials synthesis and application. Silk fibroin was modified with glycidyl methacrylate (GMA) to obtain silk fibroin methacryloyl (SFMA), while histidine was functionalized with acryloyl chloride to generate histidine methacryloyl (HisMA). SFMA and HisMA were then combined and coordinated with Zn²□ ions and incorporated with nHAP to form the full-component bioactive scaffold (SFMA/HisMA/nHAP/Zn²□). The resulting material can be 3D-printed and loaded with endothelial progenitor cells (EPCs) to yield a multifunctional scaffold with antibacterial activity, pro-osteogenic and pro-angiogenic properties, favorable physicochemical characteristics, and excellent biocompatibility, making it a promising candidate for the treatment of infected bone defects.

## 1. Introduction

The clinical management of large bone defects complicated by bacterial infection represents a persistent and formidable challenge in orthopedic and reconstructive surgery[1]. These complex injuries, which frequently arise from high-energy trauma, tumor resection, revision arthroplasty or chronic osteomyelitis, are often characterized by massive bone loss, persistent inflammation, impaired vascularization, and the establishment of biofilm-forming bacteria such as *Staphylococcus aureus*[2,3]. Infected bone defects not only delay fracture union but also contribute to prolonged disability, repeated surgeries, and in severe cases, limb amputation. The incidence of post-traumatic osteomyelitis following open fractures is estimated to range between 4.5–20%, with higher rates reported in patients undergoing multiple surgical interventions [4]. These conditions impose a heavy clinical and socioeconomic burden, emphasizing the urgent need for advanced therapeutic strategies capable of simultaneously controlling infection and promoting bone regeneration. Current clinical approaches for repairing infected bone defects primarily rely on a combination of aggressive debridement, systemic or local antibiotic therapy, and bone grafting [5]. Autologous bone grafting remains the gold standard due to its osteoconductive, osteoinductive, and osteogenic properties [6]. However, it is associated with significant drawbacks including limited donor availability, donor site morbidity, and variable resorption rates [7]. Allogeneic grafts, while more abundant, carry risks of immunogenic rejection and disease transmission, and exhibit lower osteoinductivity [8]. Synthetic graft substitutes and antibiotic-loaded bone cements, such as polymethyl methacrylate (PMMA), can provide temporary mechanical support and local antimicrobial activity, but lack bioactivity and are unable to fully integrate with host bone[9]. Furthermore, the persistence of bacterial biofilms within infected defects renders conventional antibiotic therapies largely ineffective, as bacteria within biofilms can be up to 1000 times more resistant to antimicrobial agents compared with planktonic forms [10]. Collectively, these limitations underscore the need for next-generation scaffolds that go beyond passive structural support and actively orchestrate bone regeneration in an infected microenvironment.

An ideal scaffold for infected bone defect repair must satisfy a complex set of requirements that address both biological and mechanical challenges[11]. Mechanically, the scaffold must provide sufficient stability to withstand load-bearing conditions, while maintaining a porous, interconnected structure to allow for cellular infiltration and nutrient diffusion[12]. Biodegradation must be finely tuned to match the pace of new bone formation, preventing premature collapse or long-term persistence that could trigger chronic inflammation [13].

Equally critical are the biological properties of such scaffolds. Antibacterial activity is essential to suppress infection and prevent recolonization by pathogens. Strategies employed include antibiotic loading, antimicrobial peptide incorporation, or the use of inorganic ions such as silver, copper, and zinc[14]. Among these, zinc ions (Zn²□) are particularly attractive due to their relatively low cytotoxicity, broad-spectrum antibacterial efficacy, and established role in bone metabolism[15]. Beyond infection control, scaffolds must also promote osteogenesis and angiogenesis. Osteogenesis requires not only osteoconductivity, whereby the scaffold provides a template for new bone growth, but also osteoinductivity, the ability to stimulate progenitor cells to differentiate into osteoblasts[16]. Key osteogenic pathways include RUNX2 activation, BMP/Smad signaling, and Wnt/β-catenin cascades[17]. Angiogenesis, the formation of new blood vessels, is equally indispensable, as adequate vascularization ensures oxygen and nutrient delivery to regenerating tissues. Poor vascularity is a hallmark of infected defects and contributes significantly to impaired healing[18]. Therefore, scaffolds that can stimulate angiogenesis either by releasing pro-angiogenic factors or by incorporating endothelial lineage cells, offering a distinct therapeutic advantage[19].

Natural polymers have garnered increasing attention in scaffold design due to their biocompatibility and bioactivity[20]. Among them, silk fibroin (SF), derived from *Bombyx mori* cocoons, has been widely explored for bone tissue engineering owing to its excellent mechanical strength, low immunogenicity, slow biodegradation, and capacity to support cell attachment and differentiation[21]. SF has been successfully fabricated into films, hydrogels, and porous scaffolds for musculoskeletal applications[22]. However, unmodified SF lacks photocrosslinking functionality required for high-resolution 3D printing. To address this, methacrylation with glycidyl methacrylate (GMA) yields silk fibroin methacryloyl (SFMA), a photo-crosslinkable derivative that can be precisely patterned by digital light-based 3D printing technologies[23,24]. This modification allows fabrication of patient-specific, mechanically robust scaffolds with reproducible architectures. To further enhance the osteogenic potential of SFMA scaffolds, bioactive inorganic components can be incorporated. Nano-hydroxyapatite (nHAP), which closely mimics the mineral phase of native bone, is a well-established osteoconductive additive that improves both mechanical reinforcement and cell adhesion[25]. Zinc ions (Zn²□) exhibit antibacterial properties through disruption of bacterial membranes and biofilms, while also promoting osteoblast proliferation, differentiation, and mineralization through regulation of ALP activity and collagen synthesis[26–28]. Emerging evidence also suggests a role for Zn²□ in angiogenesis by stimulating endothelial cell migration and VEGF expression[29]. A unique innovation of the present work is the incorporation of methacrylated histidine (HisMA) as a dual-functional component. Histidine, with its imidazole side chain, serves as a natural chelator of Zn²□, enabling stable incorporation and controlled release of zinc ions from the scaffold[30]. Concurrently, the methacryloyl groups on HisMA allow its integration into the photocrosslinked SFMA network, thereby enhancing scaffold stability and functionality. This design strategy ensures sustained antibacterial activity and osteogenic stimulation without the burst release commonly associated with ion-loaded scaffolds.

While material-based cues are critical, the integration of cellular components offers a powerful approach to enhance scaffold functionality. Endothelial progenitor cells (EPCs), originally identified in peripheral blood and bone marrow, are a population of endothelial lineage cells capable of differentiating into mature endothelial cells and contributing to neovascularization[31]. EPCs play an essential role in postnatal vasculogenesis and are widely studied for regenerative applications due to their angiogenic and paracrine function[32]. Importantly, EPCs secrete a repertoire of cytokines and growth factors, including vascular endothelial growth factor (VEGF), stromal-derived factor-1 (SDF-1), and bone morphogenetic proteins (BMPs), which not only promote angiogenesis but also exert osteoinductive effects on mesenchymal stem cells (MSCs)[33]. Recent studies have highlighted the crosstalk between EPCs and osteoblast-lineage cells, demonstrating that EPC-derived paracrine signals can accelerate osteogenesis by enhancing RUNX2 activation and matrix mineralization[34]. In the context of infected bone defects, where vascular insufficiency and inflammation are key barriers to healing, EPC-laden scaffolds offer a unique opportunity to create a pre-vascularized, pro-regenerative niche. Unlike mature endothelial cells such as HUVECs, EPCs possess higher proliferative capacity, migratory potential, and in vivo vasculogenic activity[35]. Their incorporation into biomaterial scaffolds has been shown to accelerate angiogenesis, improve graft integration, and enhance bone regeneration[36]. However, few studies have explored EPC-loaded scaffolds in the setting of infected bone defects, leaving a critical translational gap that this study seeks to address.

In light of these challenges and opportunities, the aim of this study was to design, fabricate, and evaluate a multifunctional, 3D-printed scaffold based on SFMA/HisMA/nHAP/Zn²□ and loaded with EPCs. The scaffold was engineered to integrate mechanical stability, controlled biodegradability, antibacterial activity, osteoconductivity, and pro-angiogenic functionality (**Graphic Abstract**). We systematically assessed its physicochemical and biological properties in vitro, focusing on antibacterial efficacy, biocompatibility, and EPC–BMSC co-culture. Finally, its therapeutic potential was investigated in a clinically relevant murine femoral infected defect model, with outcomes assessed by micro-computed tomography (µCT), histological staining, and cytokine profiling. By combining a bioactive material platform with cellular therapy, this work introduces a promising strategy for the treatment of complex infected bone defects.

## 2. Materials and Methods

### 2.1 Materials

*Bombyx mori* silkworm cocoons were obtained from a local supplier. Sodium carbonate (Na□CO□), lithium bromide (LiBr, 99%), L-histidine (99%), acryloyl chloride (96%), sodium hydroxide (NaOH), nano-hydroxyapatite (nHAP, ≥97%), hydrochloric acid (HCl), acetone, and zinc sulfate (ZnSO□) were purchased from Macklin or Aladdin (Shanghai, China). Glycidyl methacrylate (GMA, 97%) and the photoinitiator lithium phenyl-2,4,6-trimethylbenzoylphosphinate (LAP, 99.5%) were obtained from Sigma-Aldrich (USA) and TCI Chemicals (Japan), respectively. Cell culture reagents including Dulbecco’s Modified Eagle Medium (DMEM), Minimum Essential Medium α (α-MEM), fetal bovine serum (FBS), penicillin–streptomycin, and trypsin–EDTA were purchased from Gibco (Thermo Fisher Scientific, USA). Mouse bone marrow stromal cells (BMSCs) were purchased from STEMCELL Technologies (Canada), endothelial progenitor cells (EPCs) were obtained from Lonza (Switzerland), and MC3T3-E1 pre-osteoblasts were obtained from ATCC (CRL-2594, USA). EPCs were cultured in EGM-2 medium (Lonza) supplemented with growth factors and 5–20% FBS, BMSCs were maintained in DMEM with 10% FBS, and MC3T3-E1 cells were cultured in α-MEM containing 10% FBS and 2 mM glutamine. Transwell culture plates (0.4 µm pore size, Corning, USA) were used for indirect co-culture experiments. *Escherichia coli* (ATCC 25922) and *Staphylococcus aureus* (ATCC 6538) were used for antibacterial assays. Staining reagents included Calcein-AM/EthD-1 live/dead assay kit (Thermo Fisher Scientific), phalloidin-TRITC and DAPI (Sigma-Aldrich), and primary antibodies against OCN, COL1, RUNX2, OPN, IL-6, IL-1β, and TNF-α (Abcam, UK).

### 2.2 Synthesis of Methacrylated Silk Fibroin (SFMA)

Silk fibroin (SF) was extracted from *Bombyx mori* cocoons by a standard degumming process. Briefly, 40 g of cocoons were cut into small pieces and boiled for 30 min in 1 L of 0.05 M sodium carbonate (Na□CO□) solution, followed by thorough rinsing with deionized water to remove sericin. The degummed SF fibers were dried in an oven for 36 h. To prepare SFMA, 10 g of dried SF was dissolved in 100 mL of 9.3 M lithium bromide (LiBr) solution at 60 °C under stirring for 1 h. The pH of the solution was adjusted to 9.0, after which 6 mL of glycidyl methacrylate (GMA, 211.5 mM) was added. The reaction mixture was stirred at 1000 rpm and maintained at 60 °C for 6 h. The product was filtered through Miracloth (Millipore) to remove insoluble impurities and dialyzed against deionized water using a dialysis membrane (MWCO 12–14 kDa, Spectrum Labs) for 7 days with frequent water changes. The purified solution was subsequently lyophilized using a freeze-dryer (SCIENTZ-10, Ningbo Xinzhi, China) to obtain SFMA, which was stored at –20 °C until further use.

### 2.3 Synthesis of Methacrylated Histidine (His-MA)

L-histidine (3 g) was dissolved in 10 mL of 0.1 M sodium hydroxide (NaOH) solution under constant stirring. The reaction was performed under light-protected conditions, and 1.6 mL of acryloyl chloride was added dropwise to the solution over several minutes. The mixture was allowed to react for 1 h at room temperature. The pH of the reaction solution was then adjusted to 3.0 using concentrated hydrochloric acid (HCl). The product was purified by washing three times with acetone (3 × 50 mL) and subsequently precipitated from the organic layer using anhydrous ethanol. The resulting precipitate was washed with ultrapure water, filtered, and lyophilized to obtain HisMA, which was stored in a desiccator until further use.

### 2.4 Fabrication of 3D-Printed Scaffolds

Composite bio-inks were prepared by dissolving SFMA and gelatin in deionized water to final concentrations of 10% and 5% (w/v), respectively, with 0.5% (w/v) lithium phenyl-2,4,6-trimethylbenzoylphosphinate (LAP, a water-soluble photoinitiator for visible-light crosslinking) added. Gelatin served as a sacrificial material to improve printability. For groups containing HisMA, 5% (w/v) HisMA was incorporated into the SFMA solution prior to mixing with gelatin. For nHAP-containing scaffolds, nano-hydroxyapatite (2% w/v) was dispersed in the SFMA/HisMA solution by ultrasonication for 30 min to ensure homogeneous distribution before addition of gelatin and LAP. The prepared bio-ink was loaded into a printing syringe, cooled to 4 °C to induce physical gelation, and extruded using a 3D bioprinter (BioScaffolder, Germany) with the syringe maintained at 25–27 °C and the build plate at 4–6 °C. Following printing, scaffolds were photo-crosslinked under 405 nm blue light for 2 min. The sacrificial gelatin was removed by incubating scaffolds in sterile water at 37 °C for 15 min. Four scaffold formulations were fabricated: (1) SFMA (10% w/v); (2) SFMA/HisMA (10%/5% w/v); (3) SFMA/HisMA/Zn²□; and (4) SFMA/HisMA/nHAP/Zn²□. For Zn²□ incorporation, scaffolds were immersed in 24 mM ZnSO□ solution for 30 min at room temperature, followed by thorough rinsing with ultrapure water to remove unbound ions. The resulting scaffolds were freeze-dried and stored at –20 °C until further use.

### 2.5 Chemical Structure Analysis

The chemical structures of silk fibroin (SF), methacrylated silk fibroin (SFMA), histidine (His), and methacrylated histidine (HisMA) were analyzed by ^1^H nuclear magnetic resonance (^1^H-NMR) spectroscopy (Bruker AVANCE III, 400 MHz) using D□O as solvent.

### 2.6 Elemental Composition (EDS)

Elemental composition and mapping were performed by SEM-EDS (Hitachi SU8010 equipped with Oxford EDX). Lyophilized specimens were mounted on carbon adhesive tabs and carbon-coated (∼10 nm) for EDS (to avoid Au/Pt peaks); separate samples gold-coated were used only for high-resolution SEM imaging. Spectra were acquired at 15 kV, working distance ∼10 mm, live time 60 s, three regions of interest (ROIs) per sample.

### 2.7 Morphology and Microstructure

The macro- and micro-architecture of 3D-printed scaffolds were observed by scanning electron microscopy (SEM, Hitachi SU8010). Low-magnification images (×40) were used to evaluate pore size and distribution, while high-magnification images (×1000) revealed detailed surface morphology.

### 2.8 Swelling Behavior

Pre-weighed dry scaffolds (W□) were immersed in phosphate-buffered saline (PBS, pH 7.4) at 37 °C. At selected intervals, samples were removed, gently blotted to remove excess water, and weighed (W□). The swelling ratio (SR) was calculated as:

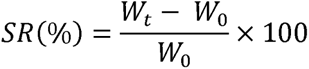

until equilibrium was reached.

### 2.9 In Vitro Degradation

Lyophilized scaffolds (W□) were incubated in PBS (pH 7.4) with or without lysozyme (1.5 µg/mL, Sigma) at 37 °C. At predetermined time points, scaffolds were collected, rinsed, freeze-dried, and re-weighed (W□). The degradation ratio was calculated as:

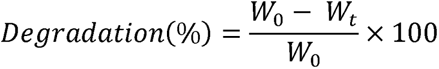

### 2.10 Rheological Properties

Rheological tests were carried out on cylindrical hydrogel samples (8 mm diameter, 1.5 mm height) using a rotational rheometer (Kinexus, Malvern, UK). Time sweep tests were conducted at 0.1% strain and 1 Hz frequency, followed by frequency sweep tests (0.1–100 rad·s□¹) at 37 °C to obtain the storage modulus (G′) and loss modulus (G″).

### 2.11 Mechanical Properties

Mechanical testing was performed using a universal testing machine (Instron 5944, USA). Dumbbell-shaped hydrogel specimens were subjected to uniaxial tensile testing at a crosshead speed of 10 mm/min. Stress–strain curves were recorded, and both tensile modulus and Young’s modulus were determined from the slope of the initial linear region. For compressive testing, cylindrical samples (6 mm diameter, 1.5 mm height) were compressed under unconfined conditions at 0.3 mm/min. All measurements were performed in triplicate, and mean values were reported.

### 2.12 Antibacterial Assessment

The antibacterial activity of scaffolds was tested against Gram-positive *Staphylococcus aureus* (ATCC 6538) and Gram-negative *Escherichia coli* (ATCC 25922). Bacteria were cultured in Luria–Bertani (LB) medium and diluted to 1 × 10□ CFU/mL. Sterile scaffold samples were incubated with the bacterial suspensions at 37 °C for 24 h. After incubation, suspensions were serially diluted, plated on LB agar, and incubated overnight. The antibacterial rate (AR) was calculated by comparing colony-forming units (CFUs) of the scaffold group (N_sample_) with the control group without scaffold (N_control_) using the formula:

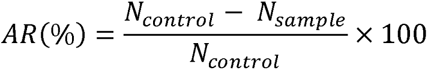

### 2.13 Biocompatibility assessment

Bone marrow stromal cells (BMSCs) were cultured in DMEM supplemented with 10% FBS and 1% penicillin–streptomycin. Endothelial progenitor cells (EPCs) were maintained in EGM-2 medium supplemented with growth factors and 5–20% FBS. Pre-osteoblast MC3T3-E1 cells were cultured in α-MEM containing 10% FBS and 2 mM L-glutamine. All cells were maintained at 37 °C in a humidified atmosphere with 5% CO□, and the medium was refreshed every 2 days. For direct culture, cells were seeded onto sterilized scaffolds placed in 24-well plates and cultured for up to 5 days. For indirect co-culture, transwell plates (0.4 µm pore size) were used: scaffolds seeded with EPCs/BMSCs/MC3T3 were placed in the lower compartment, while EPCs cells were seeded in the upper inserts in certain experimental settings. Cell viability was assessed by Calcein-AM/EthD-1 staining followed by confocal microscopy (Olympus FV3000). Cytoskeletal organization was visualized with phalloidin staining, and nuclei were counterstained with DAPI.

### 2.14 Osteogenic marker detection

To evaluate osteogenic differentiation, BMSCs were cultured on scaffolds with or without co-culture of EPCs in osteogenic induction medium (DMEM supplemented with 10% FBS, 50 µg/mL ascorbic acid, 10 mM β-glycerophosphate, and 100 nM dexamethasone). The medium was refreshed every 2 days. After 7 and 14 days of induction, alkaline phosphatase (ALP) activity was assessed using a commercial ALP staining kit. Stained samples were imaged under a bright-field microscope, and the ALP-positive ratio was calculated as the percentage of stained (ALP-positive) area relative to the total cell area using ImageJ software. Calcium deposition was evaluated by Alizarin Red S (2%, pH 4.2) staining for 30 min. Stained cultures were imaged, and the Alizarin Red-positive ratio was determined as the percentage of mineralized (red-stained) area normalized to the total culture area. For quantitative validation, bound dye was solubilized in 10% cetylpyridinium chloride and absorbance measured at 562 nm. For immunofluorescence staining, cells were fixed with 4% paraformaldehyde, permeabilized in 0.1% Triton X-100, and blocked with 5% BSA. Samples were incubated with primary antibodies against osteogenic markers (OCN, OPN, COL1A1, RUNX2) overnight at 4 °C, followed by fluorescent secondary antibodies. Nuclei were counterstained with DAPI, and confocal images were acquired using identical settings. Quantitative analysis was performed by calculating the percentage of marker-positive cells relative to the total number of nuclei. For Western blotting, proteins were extracted using RIPA buffer containing protease inhibitors, and concentrations were determined by BCA assay. Equal protein amounts were separated by SDS-PAGE and transferred to PVDF membranes. Membranes were blocked with 5% non-fat milk, incubated with primary antibodies against OCN, OPN, COL1A1, and RUNX2 overnight at 4 °C, and then with HRP-conjugated secondary antibodies. Signals were visualized with enhanced chemiluminescence (ECL), and band intensities were quantified by densitometry relative to GAPDH.

### 2.15 Proteomic analysis

To investigate the proteomic response of osteoblasts to scaffold treatment, MC3T3-E1 cells were co-cultured in a transwell system with either EPC-loaded full-component scaffolds (SFMA/HisMA/nHAP/Zn²□) or basal scaffolds (SFMA/HisMA) for 4 days. Cells were harvested, washed with PBS, and lysed in RIPA buffer containing protease and phosphatase inhibitors. Protein concentrations were determined by BCA assay, and equal amounts of protein from each sample were digested with trypsin using the filter-aided sample preparation (FASP) method. Peptides were labeled with tandem mass tag (TMT, 10-plex) reagents according to the manufacturer’s instructions and combined into a single mixture. Labeled peptides were fractionated by high-pH reverse-phase liquid chromatography and analyzed by nano-liquid chromatography tandem mass spectrometry (LC–MS/MS) using an Orbitrap Fusion Lumos mass spectrometer. Raw MS data were processed with Proteome Discoverer software against the UniProt mouse reference proteome database. Protein quantification was performed using reporter ion intensities, and differentially expressed proteins (DEPs) were identified with thresholds of |log□ fold change| > 0.3 and adjusted p-value < 0.05. Bioinformatic analyses were performed to explore functional alterations. Gene Ontology (GO) based Gene Set Enrichment Analysis (GSEA) were conducted to identify significantly enriched biological processes, cellular components, and molecular functions associated with scaffold treatment.

### 2.16 Infected Femoral Defect Model

All animal experiments were performed under protocols approved by the Institutional Animal Care and Use Committee. A murine femoral condyle defect model infected with *Staphylococcus aureus* was established. Briefly, male C57BL/6 mice (8–10 weeks old) were anesthetized with isoflurane, and a 0.8 mm diameter × 0.8 mm deep defect was drilled in the distal femoral condyle using a low-speed drill under sterile saline irrigation. Immediately after drilling, 5 µL of *S. aureus* suspension (1 × 10□ CFU/mL) was injected into the defect site to induce infection. Animals were randomized into four groups (n = 9 per group): empty defect (control), SFMA/HisMA/Zn²□ scaffold, SFMA/HisMA/nHAP/Zn²□ scaffold, and EPC-loaded SFMA/HisMA/nHAP/Zn²□ scaffold. Scaffolds were press-fitted into the defect and wounds closed with sutures. Mice were sacrificed at 1, 2, or 4 weeks post-implantation for evaluation.

### 2.17 Micro-Computed Tomography (µCT) Analysis

Femurs were harvested, fixed in 4% paraformaldehyde, and scanned by µCT (Skyscan 1176, Bruker, Belgium) at 9 µm voxel resolution, 50 kV, and 200 µA. Three-dimensional reconstructions were generated using NRecon and CTAn software. Regions of interest (ROIs) encompassing the defect site were analyzed for bone volume fraction (BV/TV) and bone mineral density (BMD). Quantitative values were averaged across three samples per group.

### 2.18 Histology and Immunohistochemistry

After µCT, femurs were decalcified in 10% EDTA (pH 7.4) for 2 weeks, embedded in paraffin, and sectioned at 5 µm thickness. Sections were stained with hematoxylin and eosin (H&E) for general morphology, Masson’s trichrome for collagen deposition, and Safranin-O/Fast Green for cartilage and scaffold remnants. For immunofluorescence and immunohistochemistry, sections were deparaffinized, rehydrated, subjected to antigen retrieval, and blocked with 5% BSA. Primary antibodies against OPN, COL1, OCN, IL-6, IL-1β, and TNF-α were incubated overnight at 4 °C, followed by HRP- or fluorophore-conjugated secondary antibodies. Nuclei were counterstained with DAPI for immunofluorescence. Images were captured using a bright-field or confocal microscope. Quantitative analysis of positive staining was performed using ImageJ software on three random fields per section.

### 2.19 Statistical Analysis

All quantitative data are expressed as mean ± standard deviation (SD). Statistical comparisons between two groups were performed using Student’s *t*-test, while multiple-group comparisons were analyzed by one-way analysis of variance (ANOVA) followed by Tukey’s post hoc test. A value of *p* < 0.05 was considered statistically significant. All analyses were conducted using GraphPad Prism software.

## 3. Results and discussion

### 3.1 Synthesis and characterization of the functional bio-ink and scaffold

To fabricate a multifunctional scaffold for infected bone defect repair, a photo-crosslinkable and bioactive hydrogel system was initially developed. The foundational organic components, silk fibroin (SF) was extracted from *Bombyx mori* cocoons and subsequently modified with glycidyl methacrylate (GMA) to produce methacrylated silk fibroin (SFMA). The successful methacrylation was confirmed by ¹H-NMR spectroscopy (**Figure 1A**). Compared to native SF, the SFMA spectrum revealed characteristic new peaks at δ = 6.12 and 5.89 ppm, attributable to the vinyl protons of the methacrylate moiety, verifying the successful modification. Similarly, L-histidine was reacted with acryloyl chloride under basic conditions to yield histidine methacryloyl (HisMA) and the functionalization of histidine was confirmed by the appearance of new signals at δ = 5.70, 6.11, and 6.21 ppm in the His-MA spectrum, corresponding to the protons of the acrylamide double bond (**Figure 1B**).These syntheses are critical in which methacrylation renders the polymers photocrosslinkable for high-fidelity 3D printing, while preserving the imidazole side chains of HisMA to act as specific chelating sites for antibacterial Zn²□ ions. Following synthesis, the polymer precursors were combined with osteoconductive nano-hydroxyapatite (nHAP) to formulate the final bio-ink. The resulting SFMA/His-MA/nHAP scaffolds were 3D printed, photocrosslinked, and subsequently immersed in a ZnSO_4_ solution to incorporate antibacterial zinc ions (Zn^2+^). Energy-dispersive X-ray spectroscopy (EDS) confirmed the stable integration of zinc throughout the scaffold. The EDS spectrum (**Figure 1C)** displays a characteristic peak for Zn, verifying that the zinc ions were successfully chelated by the imidazole groups of the histidine. The incorporated nHAP provides an osteoconductive surface mimicking bone, while the chelated Zn²□ confers not only potent antibacterial properties but also known pro-osteogenic and pro-angiogenic effects[37]. Macroscopic evaluation of the 3D-printed structures demonstrated that the incorporation of His-MA, nHAP, and Zn²□ did not compromise the printability or gross structural integrity of the scaffolds when compared to the base SFMA hydrogel (**Figure 1D**). The microstructure of the scaffolds was investigated using scanning electron microscopy (SEM) to understand how the additives influenced the internal architecture (**Figure 1E**). The control SFMA, SFMA/HisMA, and SFMA/HisMA/Zn²□ scaffolds all exhibited a relatively smooth strut surface with well-defined, regular interconnected pores. In stark contrast, the final composite scaffold (SFMA/HisMA/nHAP/Zn²□) displayed a markedly rougher and more textured strut surface, with some alterations to pore regularity. The SEM images confirm a highly interconnected porous architecture, which is critical for nutrient transport and cellular infiltration. Notably, the nHAP incorporation created a significantly rougher micro-topography on the strut surfaces. This feature is known to enhance protein adsorption and subsequent integrin-mediated cell attachment and differentiation, thus providing a more favorable microenvironment for osteogenesis[38].

**Figure 1.**
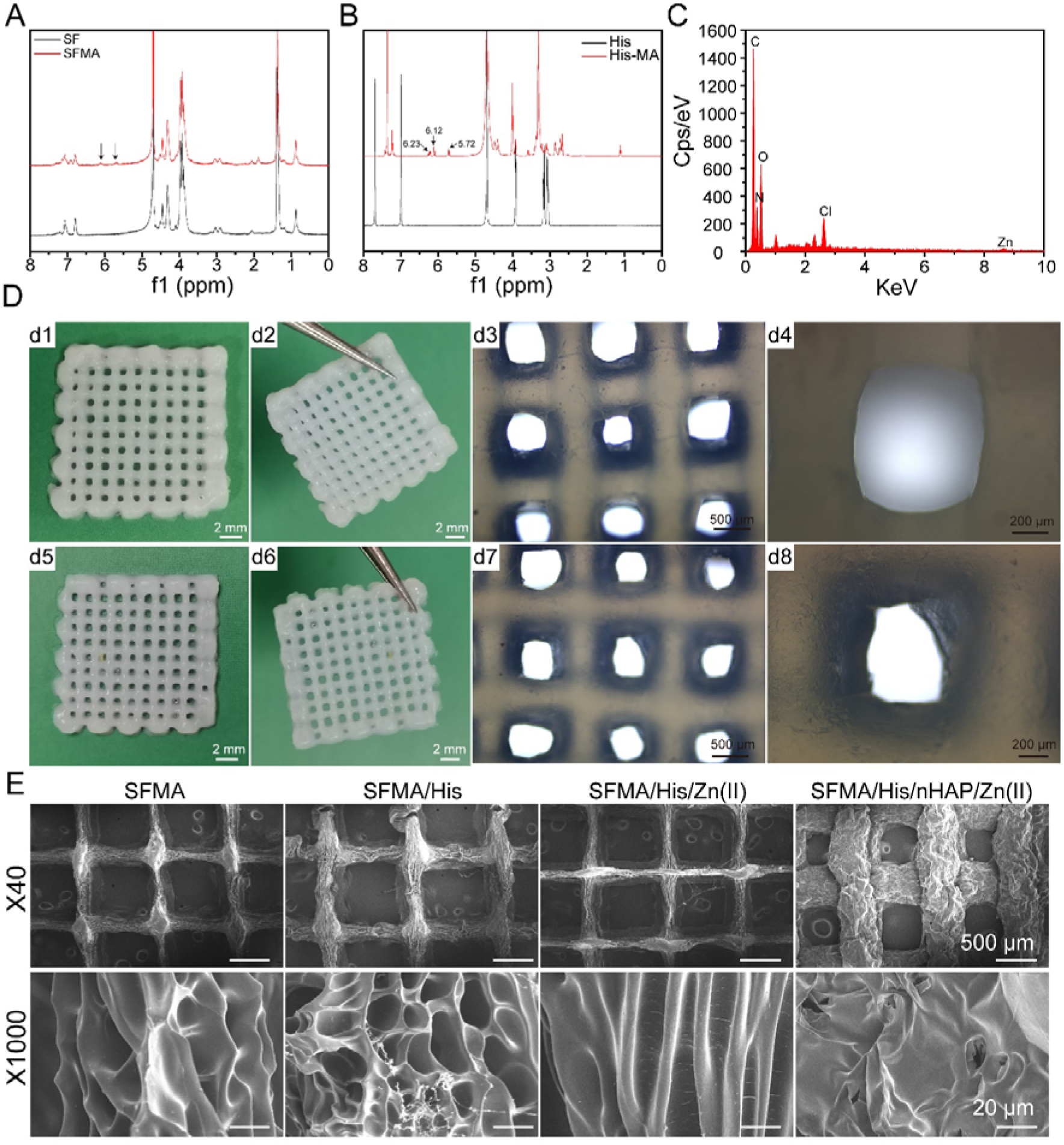
Synthesis and characterization of the functional bio-ink and scaffold. (A) ¹H NMR spectra of silk fibroin (SF, black) and silk fibroin methacryloyl (SFMA, red). (B) ¹H NMR spectra of histidine (His, black) and histidine methacryloyl (HisMA, red). (C) Energy-dispersive X-ray spectroscopy (EDS) spectrum of the SFMA/HisMA/Zn²□ scaffold. (D) Macroscopic (d1–d2, d5–d6) and microscopic (d3–d4, d7–d8) images of 3D-printed SFMA-based scaffolds. (E) Scanning electron microscopy (SEM) images of SFMA, SFMA/HisMA, SFMA/HisMA/Zn²□, and SFMA/HisMA/nHAP/Zn²□ scaffolds at low magnification (×40, scale bar = 500 μm, top row) and high magnification (×1000, scale bar = 20 μm, bottom row).

### 3.2 Physicochemical properties of SFMA/HisMA/nHAP/ Zn^2+^ scaffold

The physicochemical properties of the SFMA/HisMA/nHAP/Zn^2+^ hydrogel Scaffold were systematically evaluated to determine their suitability for *in vivo* applications as a potential bone defect implant, focusing on stability, mechanical robustness, and biodegradability. The network stability was initially tested in the swelling behavior of 3D printed scaffold (**Figure 2A**). The base SFMA scaffold exhibited the highest swelling ratio (≈35%). The incorporation of His-MA significantly reduced this to ≈23%, attributed to the formation of a secondary cross-linking network via the functionalized histidine. The subsequent chelation of Zn²□ further constrained the network, lowering the swelling ratio to ≈15%, a value that was maintained after the addition of nHAP. Crucially, the low swelling ratio of the final SFMA/HisMA/nHAP/Zn²□ scaffold is highly desirable for implantable biomaterials, as it minimizes the risk of exerting pressure on surrounding tissues post-implantation. Rheological analysis demonstrated that for all four compositions, the storage modulus (G’) consistently exceeded the loss modulus (G’’), indicative of a stable, cross-linked scaffold network with dominant solid-like elastic behavior (**Figure 2B and C**). The mechanical properties of the scaffolds were assessed under both tensile and compressive loading to ensure they could withstand the physiological environment of a bone defect. All scaffold formulations demonstrated excellent elasticity, enduring strains up to 300% without fracture (**Figure 2D**). The addition of HisMA and Zn²□ progressively increased the tensile strength compared to the SFMA control, a result of the enhanced network cross-linking. Under compression, a more nuanced behavior was observed (**Figure 2E**). The final SFMA/HisMA/nHAP/Zn²□ composite scaffold exhibited the highest compressive modulus (≈380 kPa at 55% strain), significantly greater than the SFMA control (≈330 kPa). However, while increasing stiffness, the introduction of HisMA and Zn²□ also led to increased brittleness. Notably, the incorporation of nHAP into the final composite appeared to counteract this effect, enhancing the toughness and resistance of scaffold to fracture under compression while maintaining a high compressive modulus. Finally, the biodegradability of the scaffolds, a critical feature for tissue regeneration, was evaluated over 14 days in the presence and absence of lysozyme (**Figure 2F**). As expected, all formulations were susceptible to enzymatic degradation, with lysozyme significantly accelerating mass loss compared to incubation in phosphate-buffered saline alone. The SFMA control hydrogel showed the slowest degradation kinetics. The introduction of His-MA resulted in a faster degradation profile, which was not significantly altered by the further addition of Zn²□ or nHAP. This tunable, enzyme-responsive degradation profile suggests that the scaffold can be gradually replaced by new host tissue over a clinically relevant timeframe.

**Figure 2.**
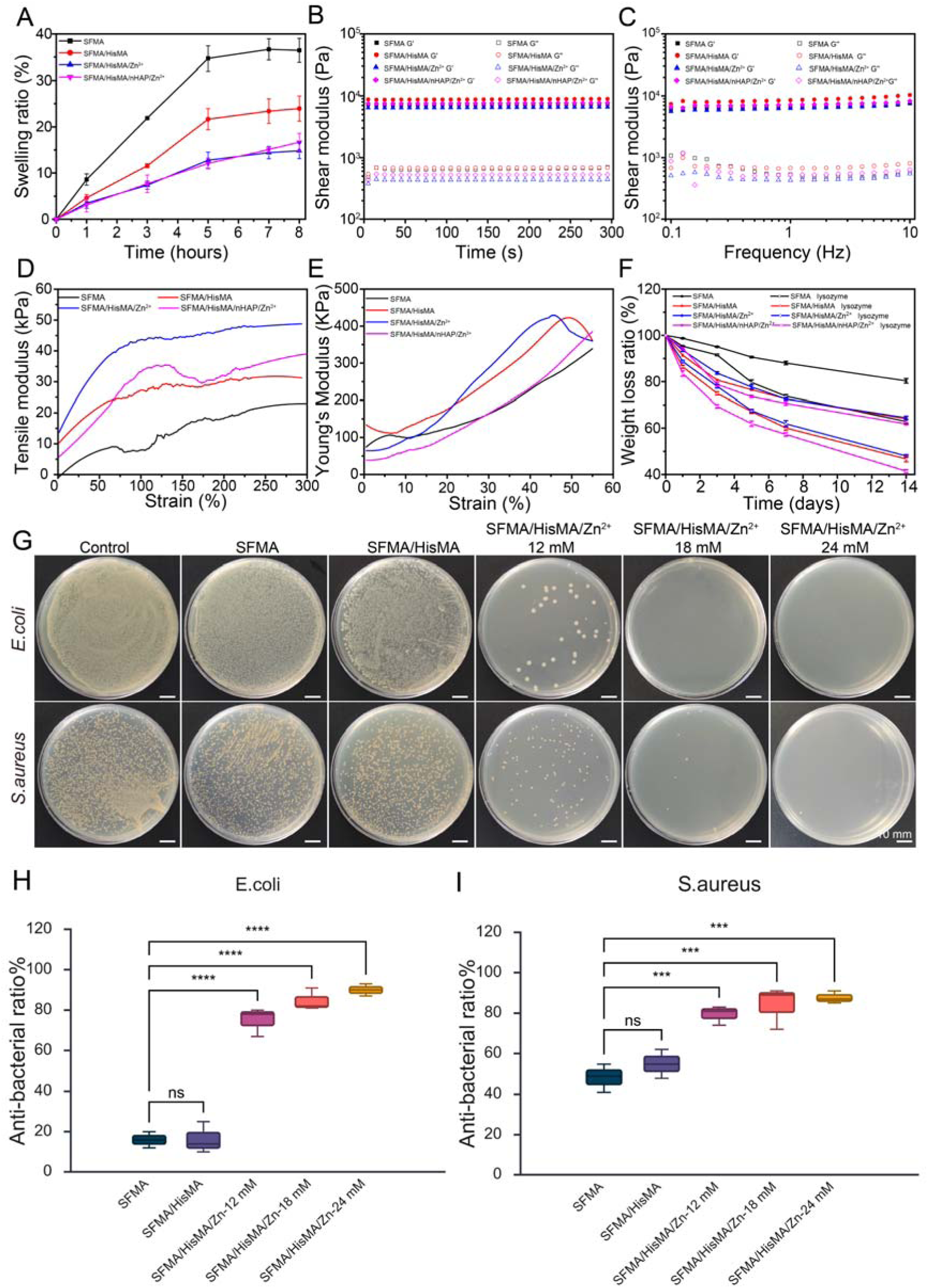
Physiochemical, mechanical and antibacterial properties of scaffolds. (A) Swelling ratios of SFMA, SFMA/HisMA, SFMA/HisMA/Zn²□, and SFMA/HisMA/nHAP/Zn²□ hydrogels measured over 8 h. (B) Time sweep rheological measurements of shear storage modulus (G′) and loss modulus (G″) for SFMA, SFMA/HisMA, SFMA/HisMA/Zn²□, and SFMA/HisMA/nHAP/Zn²□ hydrogels at 37 °C. (C) Frequency sweep rheological measurements of shear storage modulus (G′) and loss modulus (G″) for SFMA, SFMA/HisMA, SFMA/HisMA/Zn²□, and SFMA/HisMA/nHAP/Zn²□ hydrogels at 37 °C. (D) Stress–strain curves of SFMA, SFMA/HisMA, SFMA/HisMA/Zn²□, and SFMA/HisMA/nHAP/Zn²□ hydrogels. (E) Young’s modulus–strain curves of SFMA, SFMA/HisMA, SFMA/HisMA/Zn²□, and SFMA/HisMA/nHAP/Zn²□ hydrogels under uniaxial strain. (F) Weight loss ratios of SFMA, SFMA/HisMA, SFMA/HisMA/Zn²□, and SFMA/HisMA/nHAP/Zn²□ hydrogels over 14 days in PBS at 37 °C, with or without lysozyme. (G) Representative images of antibacterial assays for *E. coli* and *S. aureus*. (H, I) Quantification of antibacterial ratios against *E. coli* (H) and *S. aureus* (I). Data are presented as mean ± SD, ***P<0.0001; ****p < 0.00001 versus SFMA group, n = 3.

### 3.3. Antibacterial property and biocompatibility of SFMA/HisMA/nHAP/ Zn^2+^ Scaffold

To address the challenge of bacterial infection in bone defects, the antibacterial efficacy of the final composite scaffold was evaluated against both Gram-negative (*Escherichia coli*) and Gram-positive (*Staphylococcus aureus*) bacteria. The SFMA/HisMA/Zn²□ scaffold demonstrated potent, broad-spectrum antibacterial activity, achieving a reduction in bacterial viability of over 80% for both species (**Figure 2G and H**). Notably, this high efficiency was achieved with a Zn²□ concentration of just 18 mM. This suggests that the chelation of zinc ions by the histidine moieties not only provides a mechanism for stable zinc incorporation but also enhances its antibacterial effect at a concentration potentially lower than that required by scaffolds using unbound zinc salts[39]. Having confirmed its antibacterial activity, we then tested if the scaffold could support the growth of endothelial progenitor cells (EPCs) and bone marrow stromal cells (BMSCs) with EPCs loading (**Figure 3A**). These two cell types are critical for post trauma angiogenesis and osteogenesis respectively in vascularized bone regeneration[40]. Initially, the biocompatibility towards EPCs was confirmed. EPCs cultured on the final composite scaffold maintained a stable, well-spread morphology and exhibited higher viability at day 3 and day 5 of culture compared with SFMA/HisMA/Zn^2+^, as shown by immunofluorescence (**Figure 3B**) and live/dead staining, respectively (**Figure 3D**). To evaluate the potential of compound scaffold to support a more complex, pre-vascularized bone regeneration niche, a co-culture system was established by seeding EPCs onto the transwell above the scaffold for one day, followed by the addition of BMSCs on the scaffold. Cytoskeleton staining (**Figure 3C**) and Live/dead imaging (**Figure 3E**) over 5 days revealed excellent viability for BMSCs within the co-culture system, with significant increase in cell viability in day 5 compared with no loading of EPCs. Furthermore, cells maintained a healthy morphology, confirming that the scaffold provides a suitable microenvironment for the cohabitation and survival of both osteogenic and endothelial lineage cells.

**Figure 3.**
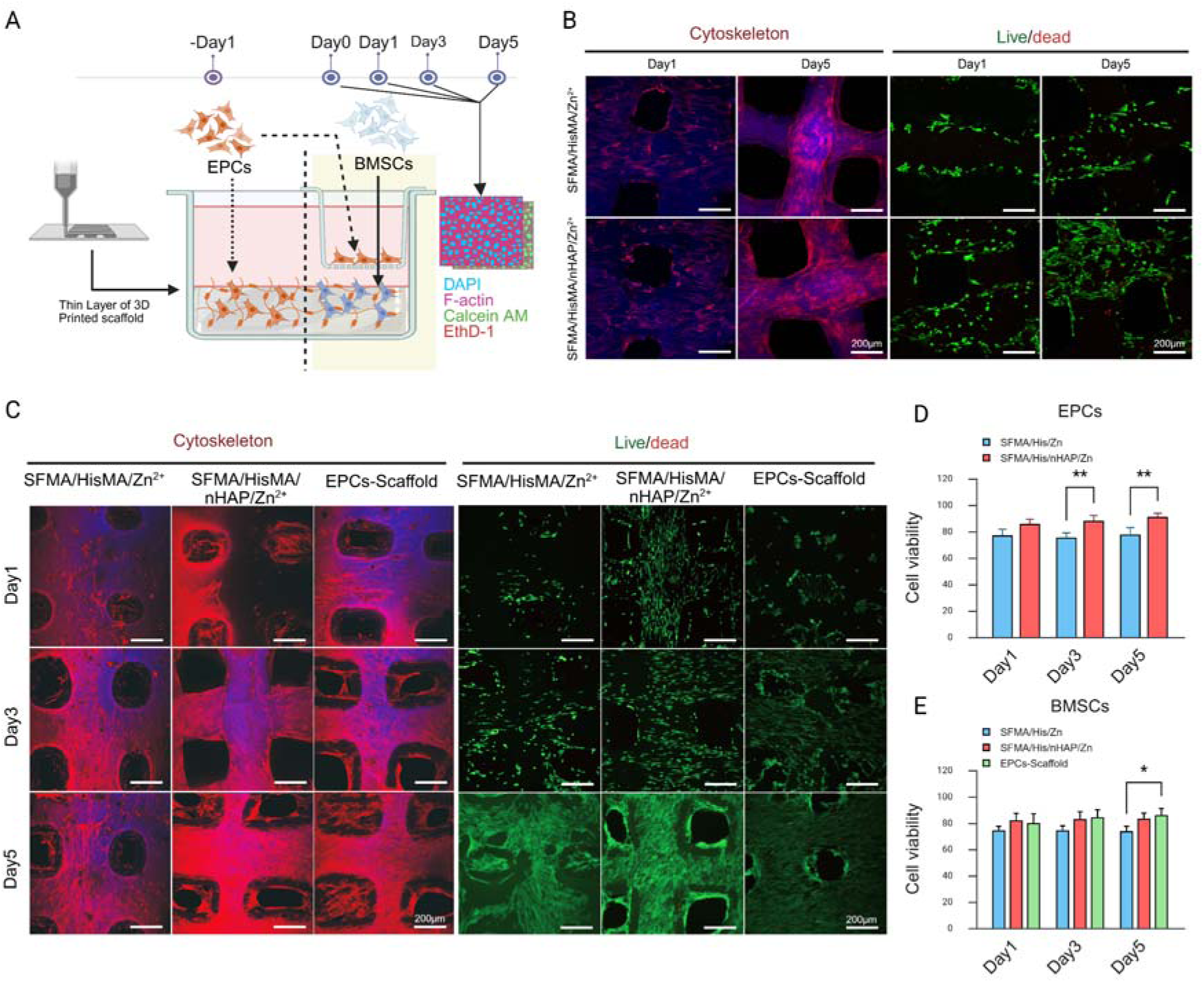
Bio-compatibility of 3D-printed functional scaffolds with EPCs and BMSCs. (A)Schematic illustration of the biocompatibility assessment. Scaffolds with different compositions were 3D printed and placed in transwell culture dishes. EPCs were seeded directly onto scaffolds for 5 days. For co-culture, BMSCs were seeded onto scaffolds while EPCs were seeded on the upper transwell insert. Cell morphology and viability were evaluated at days 1, 3, and 5 by confocal microscopy. (B) Morphology and live/dead staining of EPCs cultured on SFMA/HisMA/Zn²□ and SFMA/HisMA/nHAP/Zn²□ scaffolds. F-actin (red) and DAPI (blue) were used to stain cytoskeleton and nuclei, while Calcein AM (green) and EthD-1 (red) were applied to assess cell viability. (C) Morphology and live/dead staining of BMSCs cultured on EPC-loaded SFMA/HisMA/Zn²□, SFMA/HisMA/nHAP/Zn²□, and full-component scaffolds under co-culture conditions. (D) Quantification of EPC viability over 5 days of culture on scaffolds (n = 3, **p < 0.01 vs. SFMA/HisMA/Zn²□ group). (E). Quantification of BMCSs viability over 5 days of culture on scaffolds (n = 3, **p < 0.01 vs. SFMA/HisMA/Zn²□ group).

### 3.4 The functional scaffold triggered proteomic change on osteoblasts

As the essential bone producing cells during tissue repair of bone defect, osteoblasts have been the longstanding target for osteogenesis related therapy[41]. Therefore, the effects of this functional scaffold on osteoblasts are essential, and proteins, as the final executor of cell activity are more direct parameters to evaluate the comprehensive biological effects of 3D printed SFMA/HisMA/nHAP/ Zn^2+^ scaffold. In co-culture system, osteoblasts MC3T3 were co-cultured with EPCs loaded SFMA/HisMA/nHAP/Zn^2+^ and basal SFMA/HisMA scaffold respectively for 4 days and then osteoblasts were extracted for TMT labeled mass spectrometry for proteomic analysis (**Figure 4A**). First of all, osteoclasts grew well under both co-culture conditions but loading full component scaffold makes cell viability slightly higher than basal scaffold (**Figure 4B**). Proteomics analysis indicates that compared with basal scaffold, full component scaffold triggered 740 upregulated and 315 downregulated proteins with |logFC| >0.5(**Figure 4C and D**). Among them, a series of osteogenic, angiogenic and extracellular matrix formation genes (Flt1, Plg, Ltf, Tfrc, Wls, Serpinf1, Lum and Fbln1) were unregulated more than 2-fold while a set of pro-apoptotic signalling and metabolic stress related genes (Pak4, Pak3, Rps6ka5, Gsto1, Dbi and Pdcd5) are top down regulators. GESA based GO pathway analysis reveals that the full component scaffold orchestrates a multi-faceted, pro-regenerative proteomic regulation in osteoblasts essential for healing an infected bone defect (**Figure 4E, F and G**). The pathway enrichment results show an engagement with the acute inflammatory response, while concurrently managing the high biosynthetic demands and endoplasmic reticulum stress through enhanced mitochondrial inner membrane function. This resilient state allows for an upregulation of the endoplasmic reticulum-Golgi intermediate compartment, scaling up the cellular machinery for massive protein export. The ultimate outcome is a profound enhancement in extracellular matrix organization, evidenced by an increased capacity for collagen binding, fibronectin binding, and glycosaminoglycan binding. This coordinated activity might result in the deposition of a structured basement membrane and a robust new matrix, demonstrating the ability of scaffolds to guide osteoblasts through both immunomodulation and active tissue reconstruction.

**Figure 4.**
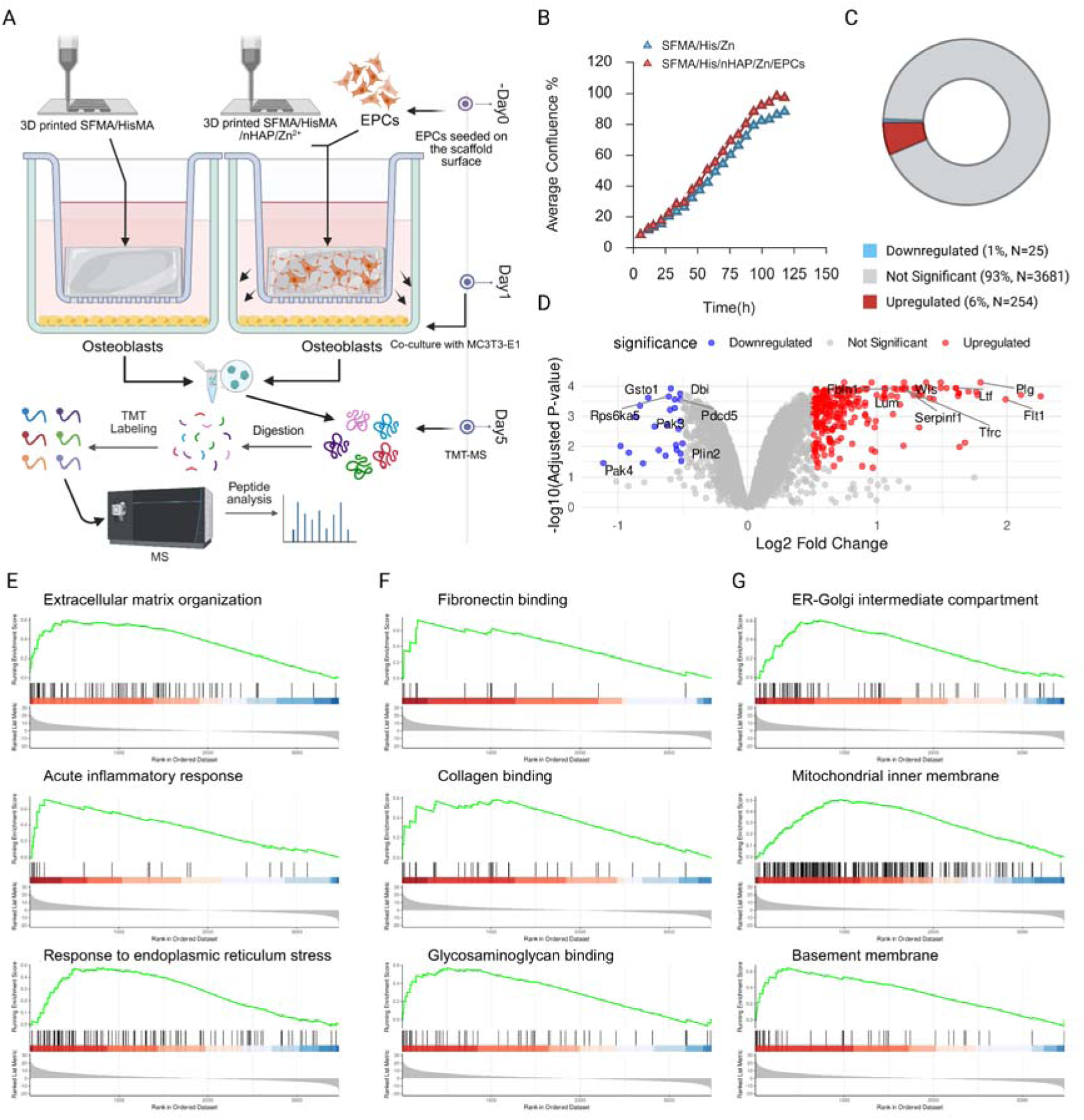
Proteomic analysis of osteoblasts co-cultured with EPC-loaded scaffolds. (A) Schematic illustration of the proteomic workflow. MC3T3 osteoblasts were co-cultured with SFMA/HisMA basal scaffolds or EPC-loaded full-component scaffolds in a transwell system. Cells were harvested at day 5 for TMT-based mass spectrometry and subsequent proteomic analysis. (B) Real-time monitoring of MC3T3 cell confluence during the co-culture period with SFMA/HisMA basal scaffolds and EPC-loaded full-component scaffolds. (C) Pie chart and (D) volcano plot of differentially expressed proteins DEPs) between control and scaffold groups. Downregulated DEPs are shown in blue, upregulated DEPs in red, and non-significant proteins in grey. (E) Gene set enrichment analysis (GSEA) of biological process terms, (F) cellular component terms, and (G) molecular function terms. The top three tissue morphogenesis-related pathways are shown as enrichment score (ES) plots for each category.

### 3.5 Scaffold-mediated osteogenesis in vitro and in vivo

Next, the primary function of the scaffold to promote osteogenic differentiation was investigated. *In vitro*, the osteogenic potential of BMSCs was assessed when co-cultured with various scaffold formulations (**Figure 5A**). Compared with SFMA control, a significant improvement in alkaline phosphatase (ALP) activity and matrix mineralization in BMSCs, as evidenced by Alizarin Red staining, was detected after adding HisMA and nHAP. More importantly, a strong synergistic effect on osteogenesis was observed after loading EPCs. (**Figure 5B, C and D**). This was corroborated at the molecular level, where immunofluorescence (**Figure 5E and F**) and immunoblotting analysis (**Figure 5G and H**) revealed a marked upregulation of key osteogenic marker proteins including osteocalcin (OCN), collagen type I (COL1), and Runt-related transcription factor 2 (RUNX2) in EPCs loaded group compared with unloaded group. The therapeutic efficacy of this scaffold was ultimately validated in a challenging *in vivo* model of a *Staphylococcus aureus*-infected femur defect in mice (**Figure 6A**). Micro-computed tomography (µCT) analysis at 4 weeks post-implantation revealed that SFMA/His-MA/Zn²□, SFMA/His-MA/nHAP/Zn²□ and EPCs loaded scaffold significantly promoted new bone formation compared to the empty defect control, evidenced by a greater BMD of femur. However, only scaffolds loaded with EPCs significantly increased the bone volume to total volume ratio (BV/TV) compared with the control group (**Figure 6B and C**). Histological analyses using H&E, Masson, and Safranin-O/Fast Green staining revealed residual undegraded scaffold material in the SFMA/His-MA/Zn²□ group, whereas substantially less residual scaffold was observed when nHAP was incorporated after 4 weeks. In contrast to scaffolds without EPCs, those loaded with EPCs exhibited more extensive and well-organized trabecular bone formation, with the scaffold material almost completely resorbed within the defect site **(Figure 6D**). These findings highlight the critical role of angiogenic stimulation in accelerating scaffold degradation and enhancing bone regeneration. Immunostaining of femoral defect sections at weeks 1, 2, and 4 demonstrated that scaffold implantation differentially modulated the inflammatory microenvironment (**Figure 6E and F**). In the control group, IL-6 expression progressively increased, consistent with unresolved inflammation, while scaffold-treated groups markedly reduced IL-6 at all timepoints. IL-1β was strongly induced by the SFMA/HisMA/Zn²□ scaffold, peaking at week 2 and remaining above control at week 4, whereas the SFMA/HisMA/nHAP/Zn²□ scaffold followed a control-like pattern but declined below control by week 4. The EPC-loaded full-component scaffold maintained IL-1β near control at early stages, with only a modest rise at week 4. TNF-α showed a sharp increase at week 1 in all scaffold groups. By week 4, only the EPC-loaded scaffold suppressed TNF-α below control, while the other scaffolds remained elevated. Osteogenic markers showed complementary trends. OPN remained stable in controls but was elevated in all scaffold groups, with the EPC-loaded scaffold showing the highest levels throughout. COL1 was transiently increased in controls at week 2 before declining, whereas all scaffolds significantly enhanced COL1 at week 1 and 2, with the EPC-loaded scaffold highest, and all groups converging by week 4. OCN decreased steadily in controls but was enhanced by scaffolds at week 1; at week 2, only the EPC-loaded scaffold maintained elevated OCN, while by week 4 both nHAP-containing and EPC-loaded scaffolds exceeded control. Together, these findings demonstrate that while all scaffolds attenuate IL-6–driven chronic inflammation and promote osteogenesis, only the EPC-loaded full-component scaffold achieves balanced immune regulation, which limit excessive IL-1β, resolve TNF-α, and sustains OPN, COL1, and OCN expression providing the most favorable environment for bone regeneration under infectious conditions.

**Figure 5.**
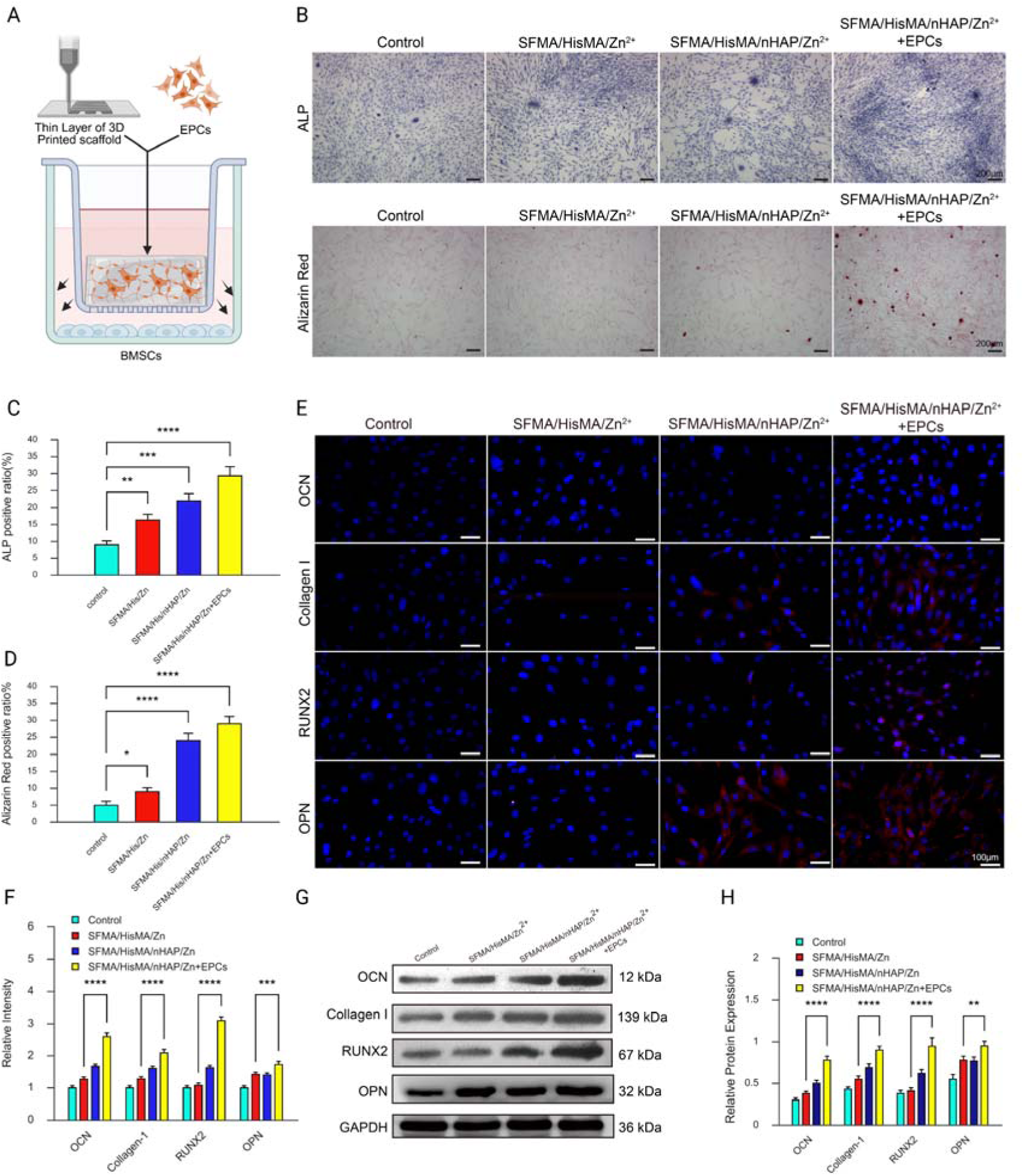
Effects of 3D-printed functional scaffolds on osteogenesis. (A) Schematic illustration of the co-culture system, in which BMSCs were cultured with EPC-loaded 3D-printed scaffolds in a transwell setup. (B) Representative images of alkaline phosphatase (ALP) and Alizarin Red staining of BMSCs under different scaffold treatments. (C, D) Quantification of ALP-positive ratio (C) and Alizarin Red-positive ratio (D) in BMSCs. (E) Immunofluorescence staining of osteocalcin (OCN), collagen I (COL1), RUNX2, and osteopontin (OPN) in BMSCs. (F) Quantification of fluorescence intensity for OCN, COL1, RUNX2, and OPN. (G) Immunoblotting of OCN, COL1, RUNX2, and OPN in BMSCs following scaffold treatment. (H) Densitometric quantification of protein expression from immunoblotting results (n = 3, **p < 0.01, ****p < 0.0001 vs. SFMA/HisMA/Zn²□ group).

**Figure 6.**
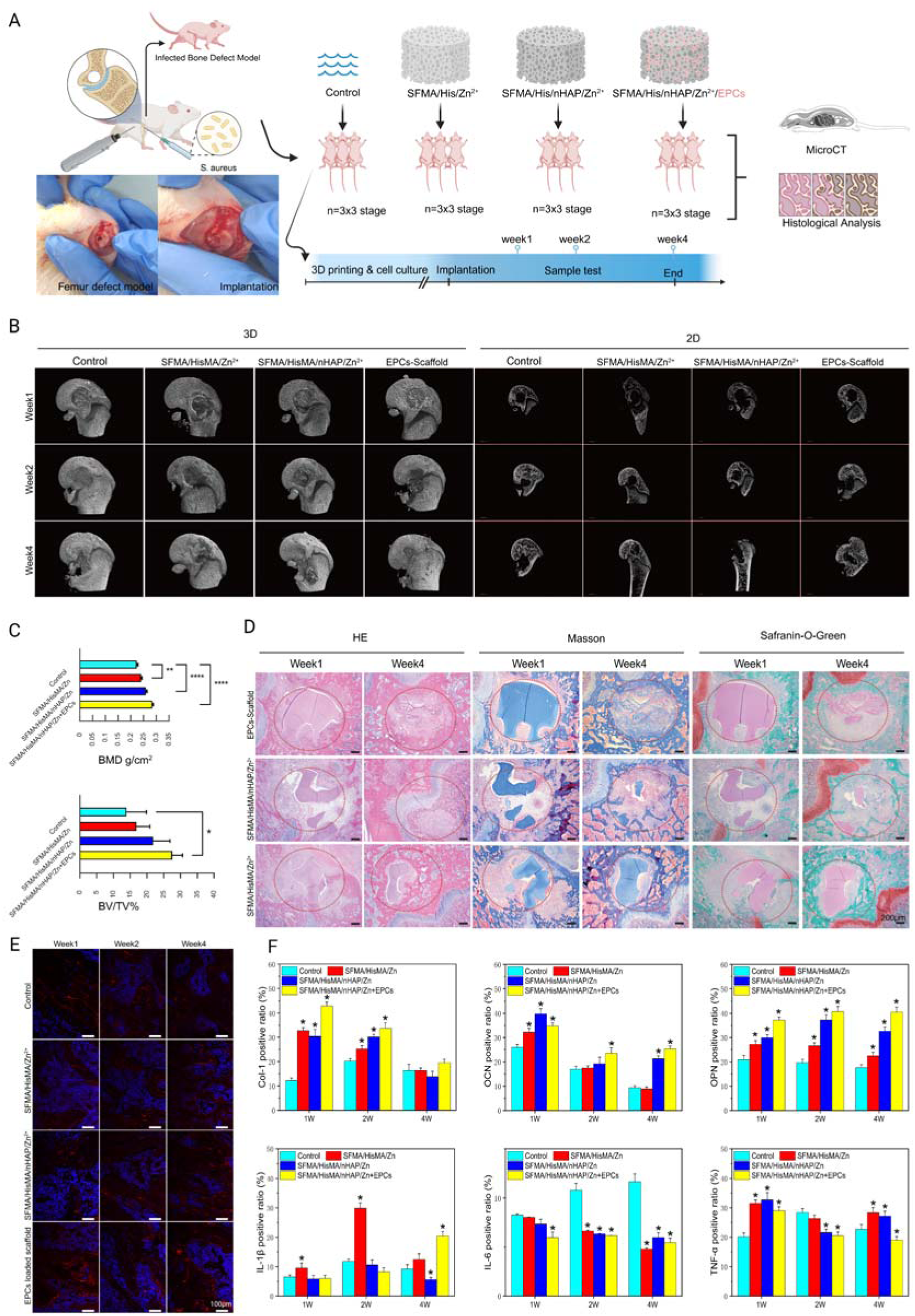
Scaffold-mediated effects in an infected bone defect model. (A) Schematic illustration of the animal model. A 3 mm diameter, 3 mm deep defect was created in the femoral condyle of mice and inoculated with S. aureus. Mice (n = 9 per group) were divided into four groups: Control, SFMA/HisMA/Zn²□, SFMA/HisMA/nHAP/Zn²□, and EPC-loaded SFMA/HisMA/nHAP/Zn²□ scaffolds. Animals were sacrificed at weeks 1, 2, and 4 for micro-CT and histological analysis. (B) Representative 2D images and 3D reconstructions from micro-CT analysis of femoral condyles in each group. (C) Quantification of bone volume fraction (BV/TV, %) and bone mineral density (BMD, g/cm²) at week 4 (n = 3, *p < 0.05, **p < 0.01, ****p < 0.0001). (D) Representative H&E, Masson’s trichrome, and Safranin-O/Fast Green staining of femoral condyle tissues at weeks 1 and 4. (E) Immunofluorescence staining of OPN in femoral condyle tissues at weeks 1, 2, and 4. (F) Quantification of immunofluorescence and immunohistochemistry results for OPN, COL1, OCN, IL-6, IL-1β, and TNF-α (n = 3, *p < 0.05 vs. control).

## 4. Conclusion

This study presents an effective regenerative strategy that integrates targeted cellular therapy with a multifunctional biomaterial to address the challenge of infected bone defects. The key innovation lies in the development of a 3D-printable silk fibroin based scaffold incorporating a histidine-methacrylate linker, enabling stable incorporation of osteoconductive nHAP and antibacterial Zn²□. When combined with EPCs, this platform established a pro-angiogenic, pro-osteogenic, and antimicrobial microenvironment. In vitro, EPC-laden scaffolds markedly enhanced the osteogenic differentiation of co-cultured bone marrow stromal cells. In vivo, this integrated system promoted robust bone regeneration and effective immune modulation in a murine femoral defect model under infectious conditions. Together, these findings demonstrate that coupling a pro-angiogenic cell source with a bioactive scaffold provides a clinically relevant platform capable of actively orchestrating bone repair, offering significant potential for treating complex orthopaedic injuries.

## CRediT authorship contribution statement

**Yunliang Zhu**: Investigation, Methodology, Formal analysis.**Jinsen Lu**: Conceptualization, Investigation, Data curation, Supervision, Writing – original draft, Writing – review & editing. **Kai Xie**: Methodology, Validation. **Shengxue Hu**: Data curation, Visualization. **Jinke Chang**: Formal analysis, Visualization. **Shiyuan Fang**: Funding acquisition, Resources. **Jiazhao Yang**: Methodology, Funding acquisition, Project administration.

## Compliance with Ethics Requirements

The animal study protocol was approved by the Institutional Animal Care and Use Committee (IACUC) of Anhui Provincial Hospital (Approval No. 2021-N(A)-200). All procedures involving live animals were conducted in strict compliance with the institutional and national guidelines for the care and use of laboratory animals.

## Acknowledgments

The authors would like to thank BIOPROFILE biotech for their assistance with the TMT-Mass Spectrometry data analysis.

